# What if wildlife health surveillance was not just for veterinarians? - Opportunistic use of population monitoring by camera traps for syndromic surveillance of the Eurasian lynx (*Lynx lynx*)

**DOI:** 10.1101/2025.05.26.656112

**Authors:** Laura Lenglin, Delphine Chenesseau, Louison Blin, Suzanne Bastian, Stéphanie Borel, Marie-Pierre Ryser-Degiorgis, Fridolin Zimmermann, Nicolas Toulet, Karin Lemberger, Philippe Gourlay, Anouk Decors

## Abstract

Fifty years after the reintroduction of the Eurasian lynx *Lynx lynx* to the Vosges and Swiss Jura mountains, the species remains in danger of extinction in France. The main threats it faces are habitat fragmentation, high anthropogenic mortality (mainly vehicle collision but also illegal killing) and low genetic diversity, but little is known about the importance of disease. Camera trapping surveillance provides crucial information for understanding population health status but includes many biases. To be more efficient, surveillance must combine several modalities. This work first presents the development of a protocol for clinical data collection by camera trapping, intended for biologists in charge of the population’s monitoring. This method was then applied to the 3574 events, i.e. 270 identified individuals, present between 1997 and 2020 in the lynx photo identification database of the Wolf-Lynx Network (Réseau Loup-Lynx) of the French Biodiversity Agency (Office Français de la Biodiversité). Seventeen percent of the lynx studied showed at least one change indicative of disease. The most common changes were skin disorders. The others were body condition, ocular or locomotor changes, lack of auriculas and rarely respiratory, behavioral, or digestive troubles. The latter changes were concentrated at the end of the study period, which also corresponded to the period with the most camera traps and therefore the most data. Finally, these observations complemented postmortem surveillance of the French lynx population carried out by the French network of epidemiological surveillance (Réseau SAGIR).

## 1. Introduction

One in four mammal species is threatened with extinction (IUCN, 2023). The Eurasian lynx *Lynx lynx*, is considered Endangered in metropolitan France (IUCN, 2023).

The role of disease in the disappearance of species is poorly studied or considered by ecologists and biologists, especially when the effects are not easy to detect, e.g. if they are sublethal or sporadic in space and time. It could nevertheless represent a far from negligible factor, depending on the species (Preece, 2017). On one hand, medium-sized cats are long-lived and have slow population dynamics. On the other hand, they are susceptible to many domestic animal diseases (e.g. feline immunodeficiency virus, feline leukemia, feline coronavirus and parvovirus, canine distemper virus and Aujeszky’s virus (Ryser-Degiorgis, 2021; Nájera, 2021)), especially in the context of a growing interface between wild and domestic animals due to the rising proportion of anthropized land. With increased infectious and possibly toxic exposure and considering the low genetic diversity of the species, it is essential for each country concerned to acquire a system for the detection of lynx diseases.

For the moment, France relies on carcass-based surveillance, which has many biases. The number of lynx that died of non-traumatic causes is probably underestimated (only 7% for the French population, according to Lena, 2020). Indeed, roadside carcasses are more likely to be found than lynx carcasses that died in remote and sparsely populated areas. Schmidt-Posthaus et al. (2002) showed that 18% of lynx found dead opportunistically were suffering from infectious diseases, but when search of lynx carcasses was facilitated using VHF collars, this proportion reached 40%. Moreover, certain organs, such as the eyes, deteriorate rapidly post mortem, making it difficult to observe lesions during necropsy. Carcass-based surveillance should thus be combined with other means of disease detection.

Camera traps (CT) have been used for several decades to estimate population size and dynamics. In the last decade, new applications in wildlife research have been developed (Rovero et al., 2013). Several studies mention the use of data collected by CT for epidemiological surveillance (Oleaga et al., 2011; Thalmann et al., 2014; Carriondo-Sanchez et al., 2017; Brewster et al., 2017; Charrier et al, 2019; Pisano et al., 2019; Muneza et al., 2019; Lacroux et al., 2019, Murray et al, 2021). However, they are still rare and mainly focused on the monitoring of cutaneous diseases that are easy to observe, such as mange in wolves (Charrier et al., 2019) or foxes (Pisano et al., 2019). The aim of our work was i) to develop a clinical observation protocol, easy to use by biologists in charge of monitoring, who can suspect pathological disorders while screening images from CTs before submitting them for final veterinary validation, ii) to apply this protocol to a dataset of images collected between 1997 and 2020 and to implement syndromic surveillance iii) compare those results with data from the SAGIR post mortem surveillance to explore how these surveillance methods complete each other.

## 2. This paper was written in parallel with Blin et al. (bioRXiv preprint, 2025)which deals with the influence of image quality on the detection of pathological changes, on the same dataset. Aspects of image quality will thus not be discussed in the present paperMaterials and methods

### 2.1. Data set

The Wolf-Lynx Network, WLN (*Fr*: Réseau Loup-Lynx, RLL) managed by the French Biodiversity Agency (Fr: Office français de la biodiversité, OFB), is dedicated since 1997 to monitoring the lynx population in France. Since 2010 it relies on systematic photo-identification by camera trapping (with the retrospective use of historical images dating back to 1997). All the images and videos used for our study came from the database dedicated to lynx photo-identification in three kinds of opportunistic detection setups: “on trail” (a paired CT set on the edge of a trail used by a lynx; 85% of recorded events, with a median of one image per event), “prey” (CT facing a carcass of lynx prey that was discovered by chance; 13% of recorded events, with a median of 8 images per event) and “direct observation” (photograph taken by a person in close proximity or even direct contact with a lynx; 2% of recorded events, with a median of two images per event). Depending on the setup, those data were collected by various correspondents such as government agents or private volunteers (shepherds, hunters, naturalists, etc.) with various observation pressures. Intensive monitoring was specifically programmed between 2011 and 2015, where paired cameras were used on each side of the trails. The images were all geolocated on a map or with GPS coordinates. The dates and times of detection were also included. All data were validated by the WLN.

A photographic “event” corresponds to all the photos or videos of an individual (or a mother with her cubs) taken at a given location, over a 24-hour period.

### 2.2. Identification of the individuals

All lynx images and videos sent by the RLL correspondents were run through an individual identification software program called Extract Compare (Conservation research LDT, 2010). It compared new lynx images with those stored in the database. When the quality of the image was sufficient, the software was able to identify individuals by comparing the fur spot pattern of forelimb, flanks and thighs with those of all the lynx recorded in the database (Figure 1). We only kept events for which the individuals were fully identified (i.e. their two flanks were known and associated with the same identifier) to avoid possible double counts of individuals.

**Fig. 1.**
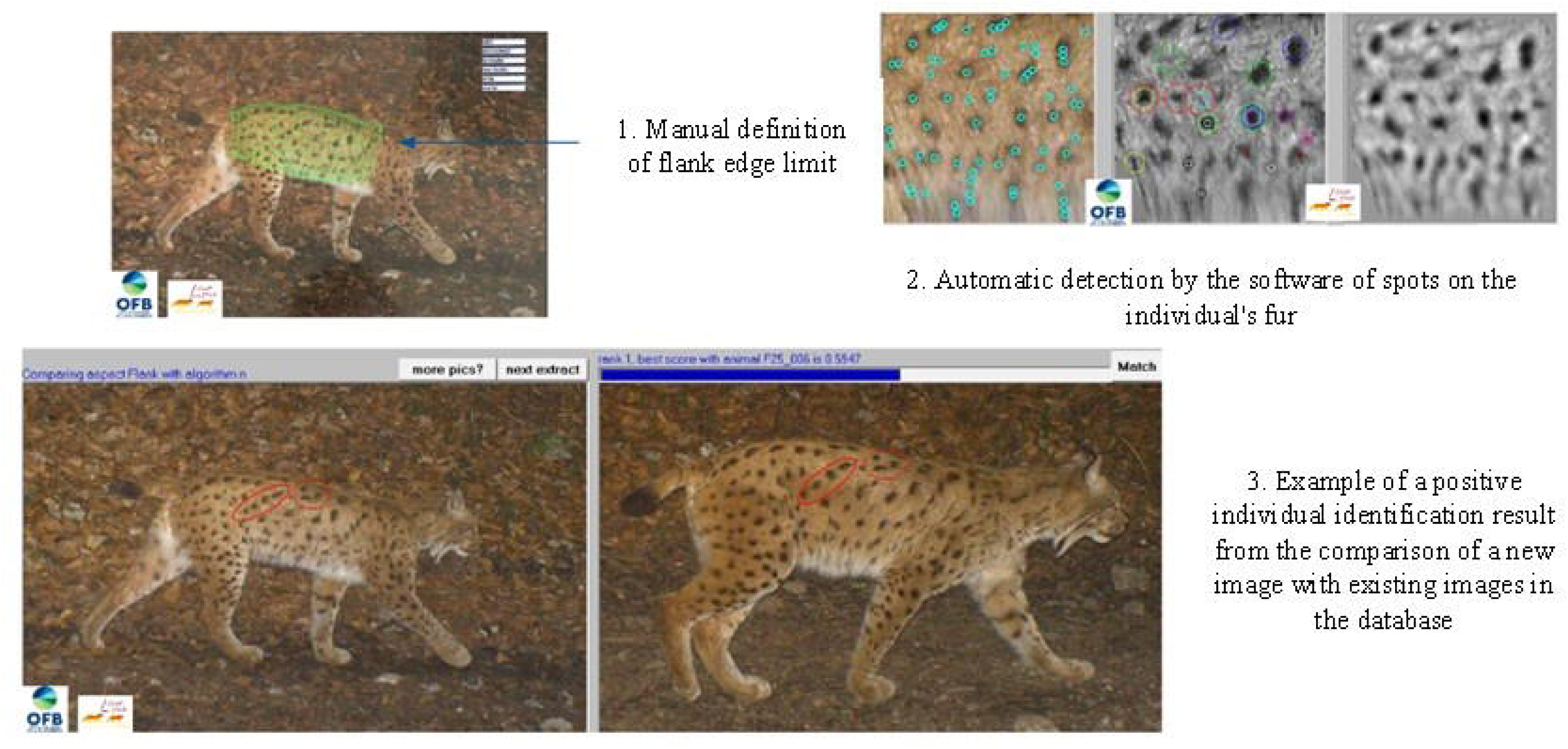
Steps to identify an individual using Extract Compare software (Individual F25_006 monitored from 2005 to 2017)

### 2.3. Clinical observation protocol and level of confidence

To limit interpretation biases and facilitate data entry and analysis, we standardized the protocol of clinical observation applied to each lynx image.

Syndromic surveillance is not based on laboratory-confirmed diagnoses but on non-specific health indicators, including clinical signs. So all images with suspected clinical changes independent of any medical or biological interpretation were sorted by two veterinary students without wildlife disease experience and then checked by a veterinarian with expertise in wildlife disease epidemiology and necropsy. The level of confidence of an event was classified into three modalities:

- ***Confirmed*** (***C***): if a clinical change is detected univocally by the expert in at least one image of the event.
- ***Doubtful*** (***D***): if a clinical change appears to be detected by the expert in at least one image of the event, but there is still some doubt and no change is detected on the other images.
- ***No Visible Change of diagnostic value*** (***NVC***): if no clinical change is detected on any of the images of the event.

Once all the lynx images were observed, we chose the most relevant images to build two atlases to help biologists standardize their observations in the future: i) a physiological lynx atlas, ii/ a pathological lynx atlas detailing which signs could be seen on a photograph or a video.

Each event recorded was characterized by: date, location, lynx identification number, circumstances of detection, presence or absence of clinical changes, category of clinical changes, characterization of clinical changes, level of confidence of the event (C/D/NVC). Any visible lesion, regardless of its severity, was recorded. The terms for describing changes were chosen to be easily appropriated by non-veterinarians, but sufficiently precise to enable syndromic surveillance. The categories of changes were based on a guide for French veterinary pathology laboratories used since 2016 (unpublished).

### 2.4. Comparison with the existing carcass-based surveillance by the SAGIR network

To assess the appropriateness of combining different surveillance systems, we compared qualitatively the nature of the lesions and the detection moment obtained in the framework of SAGIR and our opportunistic CT surveillance.

In France, the actual lynx disease surveillance relies on the event-based disease surveillance network ‘SAGIR’, which is dedicated to epidemiological surveillance of abnormal mortality events in birds and mammals (opportunistic sampling method) (Decors et al., 2022). SAGIR surveillance relies on the investigation of the cause of death through a trans-disciplinary approach, involving field clues, post-mortem examination, laboratory analyses and scientific expertise. It is based on a collaboration between the OFB, hunter’s federations and public veterinary analysis laboratories. A network of RLL volunteers of diverse profiles are coordinated at the departmental level by two technical contacts (one within the hunter’s federation, the other within the OFB). Note that one-third of the animals necropsied are observed alive (dying) beforehand. Moribund animals can also be filmed to inform veterinary expertise (Millot et al., 2017; Decors et al., 2022).

First, we compared our results with causes of death of 175 lynx necropsied by the SAGIR network between 1990 and 2019, as described by Lena et al. (2022). Then, we extracted the data from the SAGIR necropsy database and renamed the lesions with the terms defined for this study to be able to compare the results. Between 2011 and 2022, 140 lynx carcasses were managed by SAGIR. Due to intense traumatic lesions, masking certain sublethal processes, we chose to retain only necropsied individuals that were not victims of collision (i.e. 38/140).

## 3. Results

This study included 270 identified individuals corresponding to 3 574 events containing 26 828 images and 1 345 videos, collected between 1997 and 2020. The median number of lynx images per event was two. Some images that were obtained after 2020 were presented in the atlas because they provided a better illustration of specific changes, but were not considered in the data analysis from 1997 to 2020.

### 3.1. Clinical observation method

#### 3.1.1. Observation protocol

We defined a clinical observation protocol to help biologists analyze each image and video in a standardized way and detect the presence of any changes (see supplementary material). Before looking for clinical changes, we considered the setup of the capture, the season, the time of day (day *vs.* night) and the weather conditions which can influence the quality of the data and the interpretation of the image (for example, to differentiate molting from pathological hair loss). For more details, see Blin et al. (bioRXiv preprint 2025) Then the general condition of the lynx was evaluated: body condition, general aspect, and posture. An emaciated individual was characterized by visible bony protrusions (protruding shoulder, gluteal and hip tips, ribs, and spinous and/or transverse processes of the spine). Because the body condition is difficult to assess without palpation, only two modalities were defined for each individual on a given date: normal or insufficient. Any neurological or behavioral changes were also checked especially via video resources. Whenever the head was visible, eyes, nose and ears were carefully observed. Images of suspected lesions taken during the day were zoomed and contrasted to see color changes for example. Backsides were observed too, to detect any anal/vaginal secretions, amyotrophy. Videos were used to detect locomotor anomalies (lameness), as well as certain digestive, respiratory and neurological changes.

#### 3.1.2. Lynx CT atlas: physiology and changes observed

The detailed examination of the images in this database allowed us to classify them into categories of physiological and pathological visible changes. Two atlases were made from these observations, one to train the observers’ eye to recognize physiological characteristics of the species, the other to recognize and give consistent names to pathological changes.

The first atlas, (see supplementary material) presents images of individuals of different ages, sexes, and coat patterns (rosettes vs spot), to give an idea of the morphological diversity of the species. They were presented at different seasons, by day and night, taken with different CT (black and white pictures *vs.* colors) to accustom the observer’s eye to the different observation conditions and aspects of the coat. A special section is dedicated to the classification of emaciation.

In the second atlas, clinical changes types were defined. We created sub-categories to better characterize the clinical changes observed. Changes in italics had not been observed before 2020. They were however maintained in the atlas since future observation remains possible.

- Body condition change (Tbc_)
- Behavioral change (Tbehav_)
- Digestive change (Tdig_)
- *Genital change (Tgen_)*
- Locomotor change (Tloc_)
- Lack of auriculas
- *Neurological change (Tneuro_)*
- Ocular change, uni or bilateral (Toc_Ul_ or Toc_Bl)
- Respiratory change (Tresp_)
- Skin change (Tskin_)

### 3.2. Clinical changes observed in the population

#### 3.2.1. Temporal distribution of lynx with clinical changes

Out of the 270 animals identified in the database, 46 individuals (17%) showed at least one clinical change. They appeared in 3% of events (102/3574 events).

Two hundred and thirty-three individuals were recaptured at least once (*i.e.* 85%), and among them 39 individuals showed clinical changes. The median number of recaptures by individual was seven with a minimum of one and a maximum of 182. Among the recaptured lynx, there was a worsening of hair loss in 40 % of recaptured lynx with that pathology or an improvement in 20% of them. Wound healing was also observed in subsequent sightings of a same individual.

The distribution of detected individuals (with or without changes) was not homogeneous over time (Figure 2). The number of lynxes detected per year globally increased during the study period, with a minimum of 50 individuals per year, from 2011 onwards. The first lynx affected by a change was detected in 2007 and then in 2011. Thereafter, changes were recorded every year. The proportion of lynx affected rose from 0 to 10% before 2016, then varied from 6 to 21 % each year, with an average of 15.5%.

**Fig. 2.**
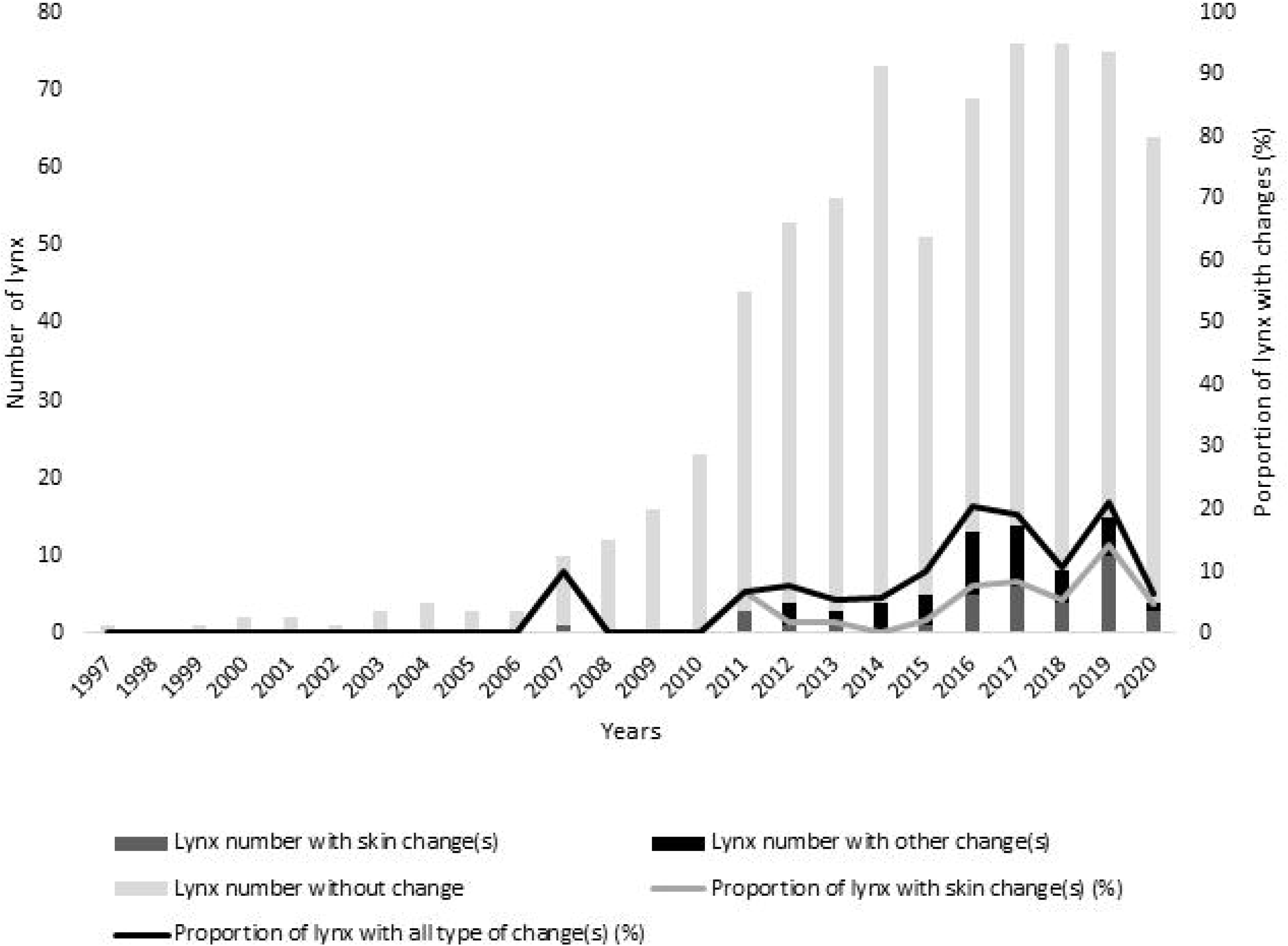
Number and proportion of individuals with clinical changes (all types combined and skin changes) in the population, between 1997 and 2020 (n=270 lynx)

#### 3.2.2. Different types of clinical changes observed

Skin changes were the most frequently observed, accounting for more than half of events with anomalies, and were identified in almost 10% of the individuals in the total population observed. Then in decreasing rank order we observed: body condition changes (4.4%), ocular changes (3.3%). Lack of auricles, locomotor, behavior, digestive and respiratory changes were rarer (Table 1). Eleven of the 46 individuals affected presented at least two different changes during their surveillance period.

**Table 1.** Number of individuals with clinical changes and proportion by type of change (T) in the affected lynx population (n=46) and in the whole population (n=270) in France between 1997 and 2020.

|  | Number of affected individuals | Proportion of lynx with this type of change among lynx affected by at least one change (%) (n=46) | Proportion of lynx with this type of change among identified lynx (%) (n=270) |
| --- | --- | --- | --- |
| **Body condition changes** | **12** | **26.1** | **4.4** |
| Distended abdomen | 1 | 2.2 | 0.4 |
| Thinness | 11 | 24.0 | 4.0 |
| **Behavior changes** (lack of leakage behaviour) | **2** | **4.3** | **0.7** |
| **Digestive changes** (vomiting) | **1** | **2.2** | **0.4** |
| **Locomotor changes** | **3** | **6.5** | **1.1** |
| Amyotrophia | 1 | 2.2 | 0.4 |
| Amputation | 1 | 2.2 | 0.4 |
| Lameness | 1 | 2.2 | 0.4 |
| **Bilateral lack of auricula** | **3** | **6.5** | **2.2** |
| **Ocular changes** | **9** | **19.6** | **3.3** |
| Color modification | 1 | 2.2 | 0.4 |
| Inflammation | 2 | 4.3 | 0.7 |
| Prolapse third eyelid (Unilateral) | 3 | 6.5 | 1.1 |
| Prolapse third eyelid (Bilateral) | 4 | 8.7 | 1.5 |
| **Respiratory changes** (nasal discharge) | **1** | **2.2** | **2.2** |
| **Skin changes** | **26** | **56.5** | **9.6** |
| Severe hair loss | 5 | 10.9 | 1.9 |
| Minimal hair loss | 5 | 10.9 | 1.9 |
| Rat tail | 10 | 21.7 | 3.7 |
| Scar | 4 | 8.7 | 1.5 |
| Deep wound | 1 | 2.1 | 0.4 |
| Superficial wound | 5 | 1.9 | 1.9 |

#### 3.2.3. Body condition changes

Twelve animals showed a degraded body condition over the study period (17 events). This category included different emaciation stages (no detailed here) and one case of abdominal distention. Stages of leanness ranged from mild emaciation to cachexia (Figure 3).

**Fig. 3.**
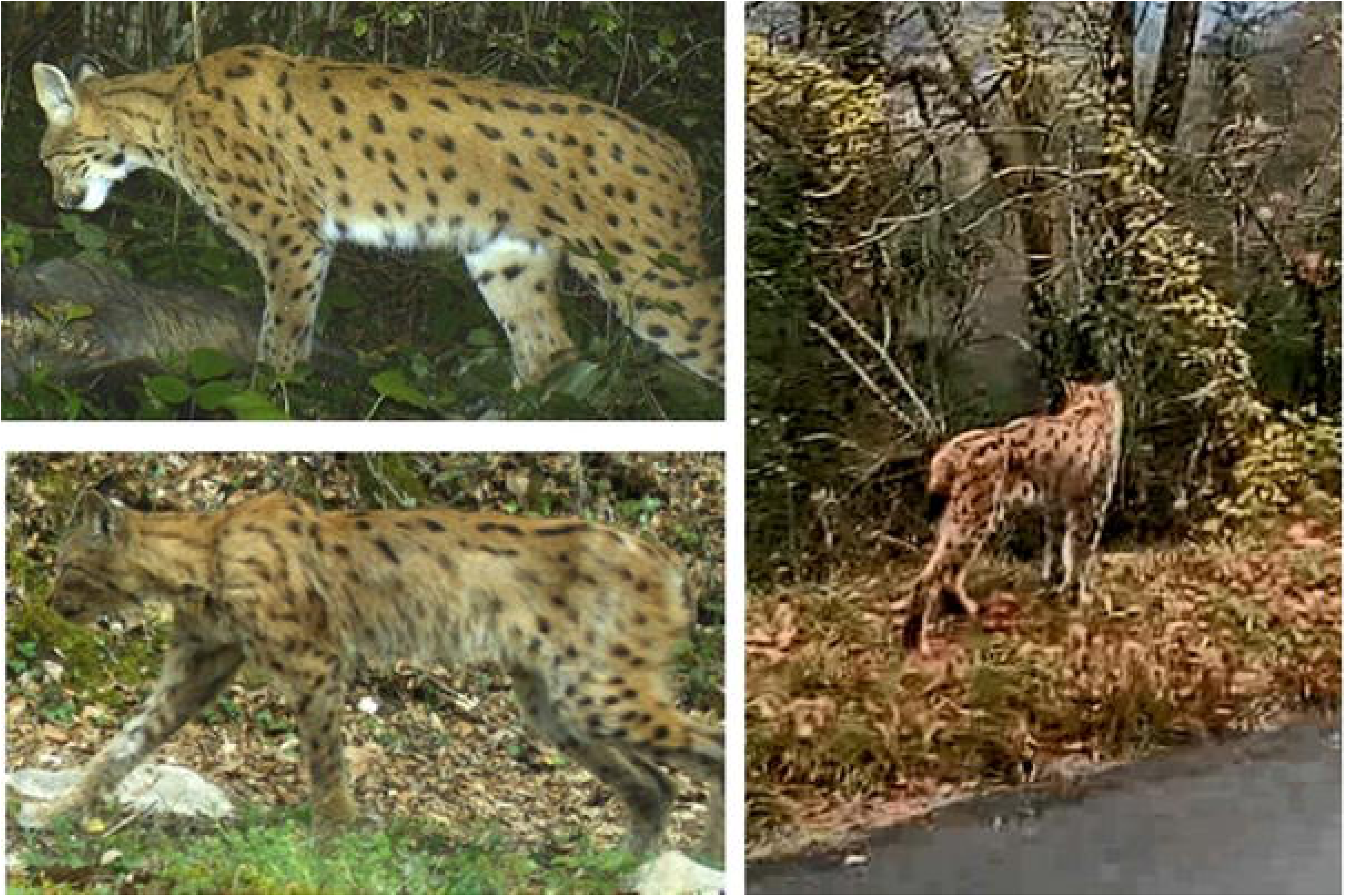
Examples of body condition observed by camera-trap on lynx – A (*on the top left)*: an individual in good body condition (©FDC 39; 08-01-2017) ; B *(at the bottom left)* : leanness characterized by the prominence of the point of the shoulder, the buttock, the hip, the vertebral spinous processes as well as the marked hollow of the flank (individual also showing a dull cot and alopecic areas on the flanks) (©FDC01; 03-21-2017) ; B *(on the right):* Very poor body condition suspected, characterized by a pronounced flank depression and a marked amyotrophy in the lumbar region and the thighs. Due to the asymmetric aspect of forehand and hindquarters, we cannot exclude an amyotrophy following a bone damage in the lumbar region (©OFB39; 11-04-2017)

Changes in body condition were observed between 2015 and 2019 and the main change was emaciation. (Table1). In 70.5 % of cases, these changes had a “confirmed” status (validated by an experienced wildlife veterinarian).

#### 3.2.4. Skin changes

Twenty-six individuals were recorded with skin changes (which corresponded to 60 events). Skin changes included hair loss of varying severity and location, wounds, and scars (Figure 4).

**Fig. 4.**
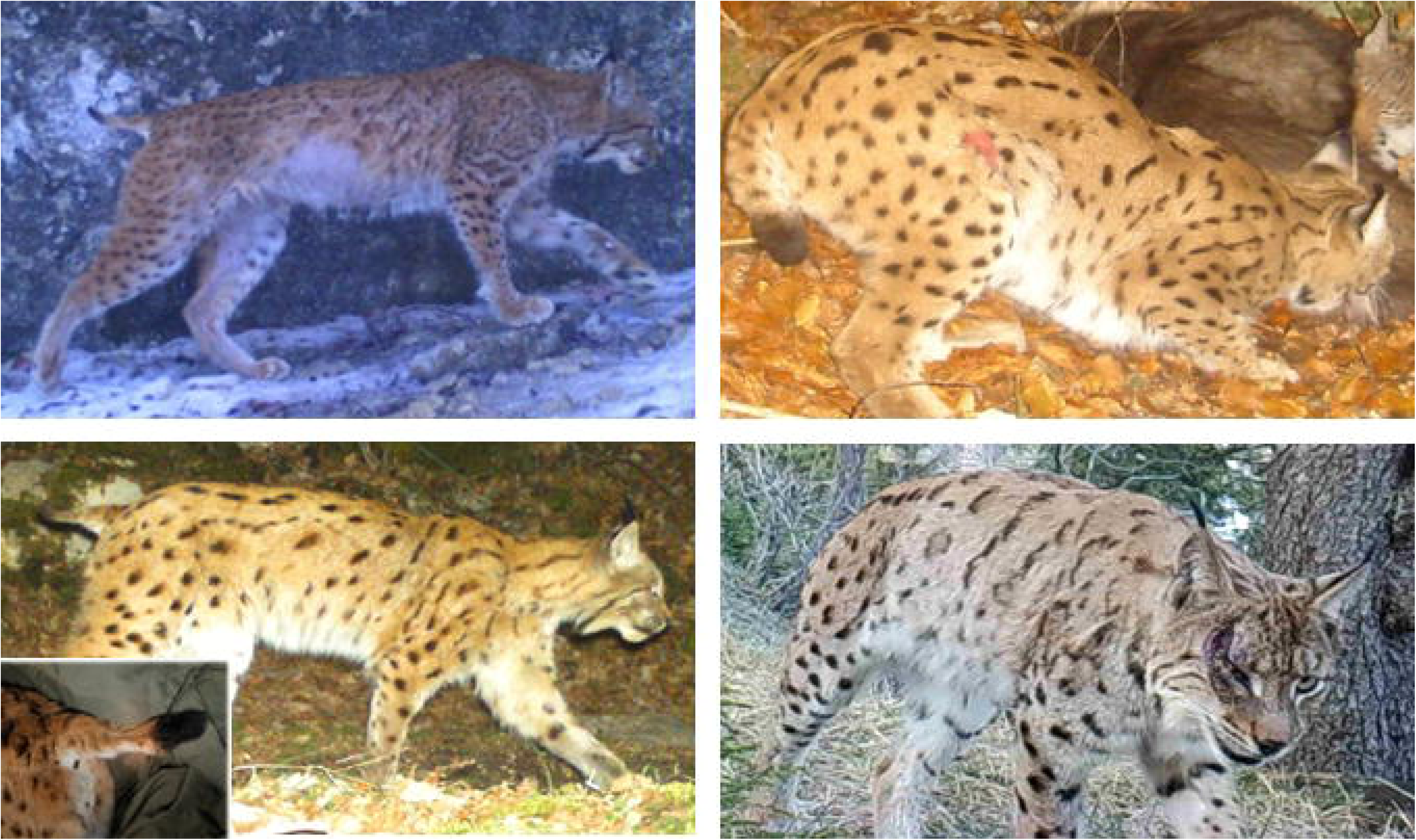
Examples of skin changes detected by camera trap on lynx – A (*on the top left)*: Severe depilation on abdomen, tail, and medial sides of limbs (©FDC25; 01/22/2021); B *(on the top right)* : Superficial wound on the right stifle (©FDC39; 10/12/2007); C (*on the bottom left, big picture*) : "Rat tail syndrome" © FDC01; 02/14/2017; D *(on the bottom left corner, small picture)* : "Rat tail syndrome", Necropsy picture: © M-P. Ryser Degiorgis); E *(on the bottom right)* Deep wound on the lateral quantum of the right eye (©JM Thevenard; 04/01/2019).

On average, skin changes were the most frequently observed changes. Generally, the proportion of skin affected individuals followed the same trend as the proportion of lynx affected by any change each year (Figure 2). But in some years, as in 2014, little or no skin changes were observed.

The most common skin changes detected were hair loss and “rat tails,” each accounting for a third of the changes observed in this category (Table 1). The level of confidence associated was 100% confirmed for severe hair loss, deep wounds, and scars. Superficial wounds were confirmed in 90% of cases. Rat tail syndrome was confirmed in 30% of suspected cases. Over the entire study period, 12.4% of identified lynx suffered from skin changes in spring (summer: 4.7%, winter: 2.9%, autumn: 2.3%). Note that for three lynx, alopecic lesions were observed cyclically in the spring (or few days before or after spring season), then disappeared and reappeared the following year at the same season (up to 3 years in a row).

#### 3.2.5. Eye changes

Eye changes affected nine individuals which corresponded to nine events. Ocular changes included conjunctival/scleral inflammations, light reflection changes, and uni- and bilateral prolapse of the third eyelid (Figure 5).

**Fig. 5.**
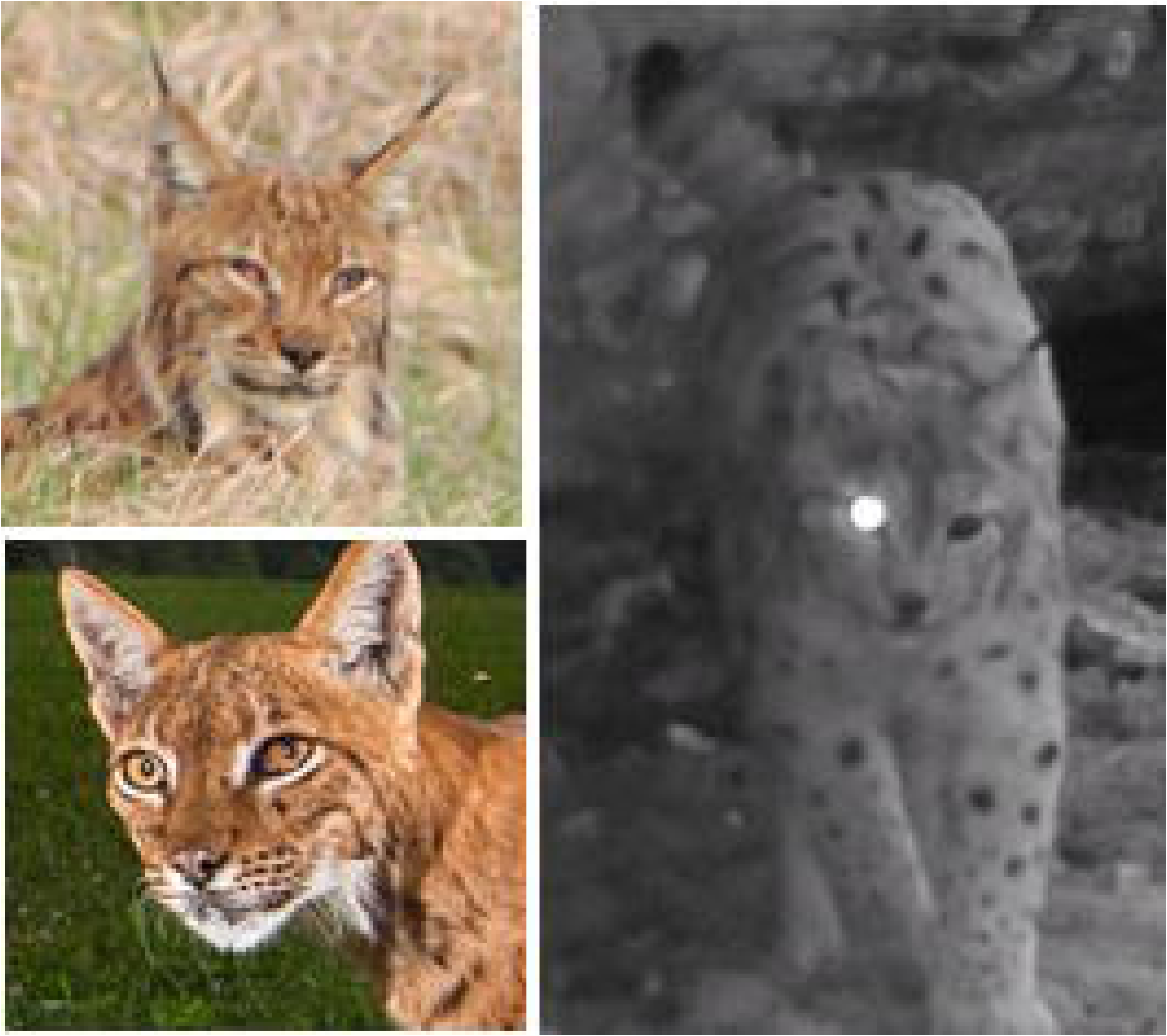
Examples of ocular specific detections by camera traps of the lynx – A (*on the top left*): Severe bilateral conjunctivitis and prolapse of the 3rd eyelid (©FDC39; 03/27/2012) B (*on the bottom left*): Dull-colored iris on the left eye with blepharospasm associated with slight prolapse of the 3rd eyelid, lesion in favour of uveitis (©Rondeau; 07/23/2013). On the right: representative screenshot of one video showing a lack of light reflection on the *tapedum lucidum* of the left eye which demonstrates the value of flash in detecting eye health anomalies (©FDC39 10/02/2022)

Eye changes were mainly observed between 2012 and 2017, only one individual was observed in 2019.

Third eyelid prolapse (uni or bilateral) accounted for 70% of ocular changes, often accompanied by ocular inflammation (on two individuals; suspicion of conjunctivitis and/or uveitis). The case of modification of flash reflection corresponded to a modification in the *tapedum lucidum* reflection detected by the camera at night with flash (which can indicate a lesion of a structure rostral to the retina). The expert veterinarian confirmed eye changes in 50% of the suspected cases but could not confirm the rest.

#### 3.2.6. Other changes

The other types of changes observed were very rare. The few locomotor changes were detected on video and varied in nature: one case of amyotrophy, one distal forelimb amputation and one case of lameness of unknown origin. Forty percent of those observations had a “confirmed” level of confidence. Bilateral absence of auricles had a level of confidence “confirmed” in all cases (Figure 6). There were two cases of behavioral disorders, one of two “confirmed”, characterized by absence of flight when a human approached at a short distance. The digestive change was represented by a lynx vomiting (doubtful observation), and the respiratory change consisted of nasal purulent discharge (confirmed observation).

**Fig. 6.**
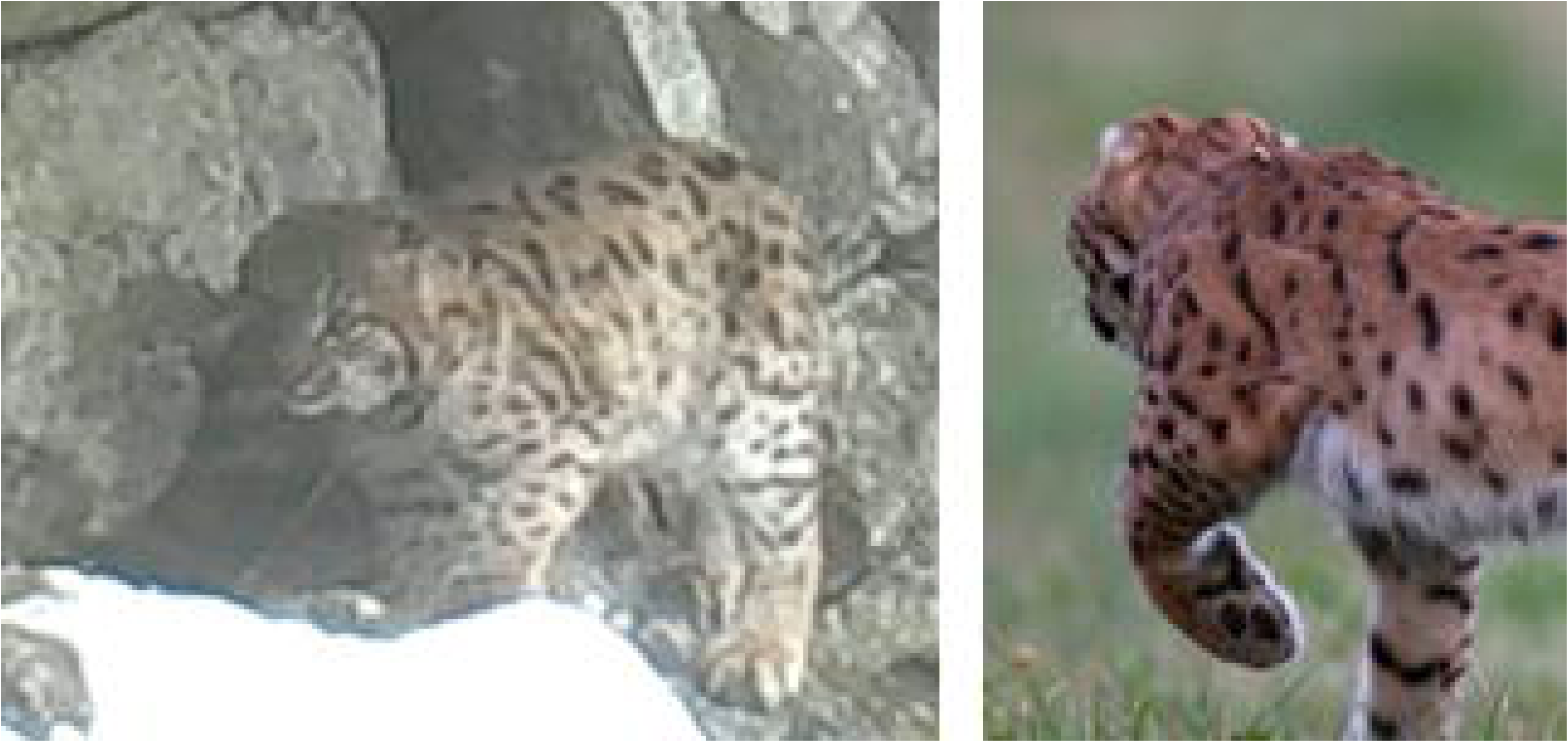
Examples of auricle lack: A (on the left): a young individual (©JM Thevenard; 11/18/2019); B (on the right): an adult (©M Balanche; 07/26/2022)

### 3.3. Contribution of the CT surveillance to global event-based surveillance

Between 2011 and 2022, out of 140 carcasses of lynx managed by SAGIR, 38 that died of non-traumatic causes were selected for a comparison with CT monitoring (Table 2).

**Table 2:**
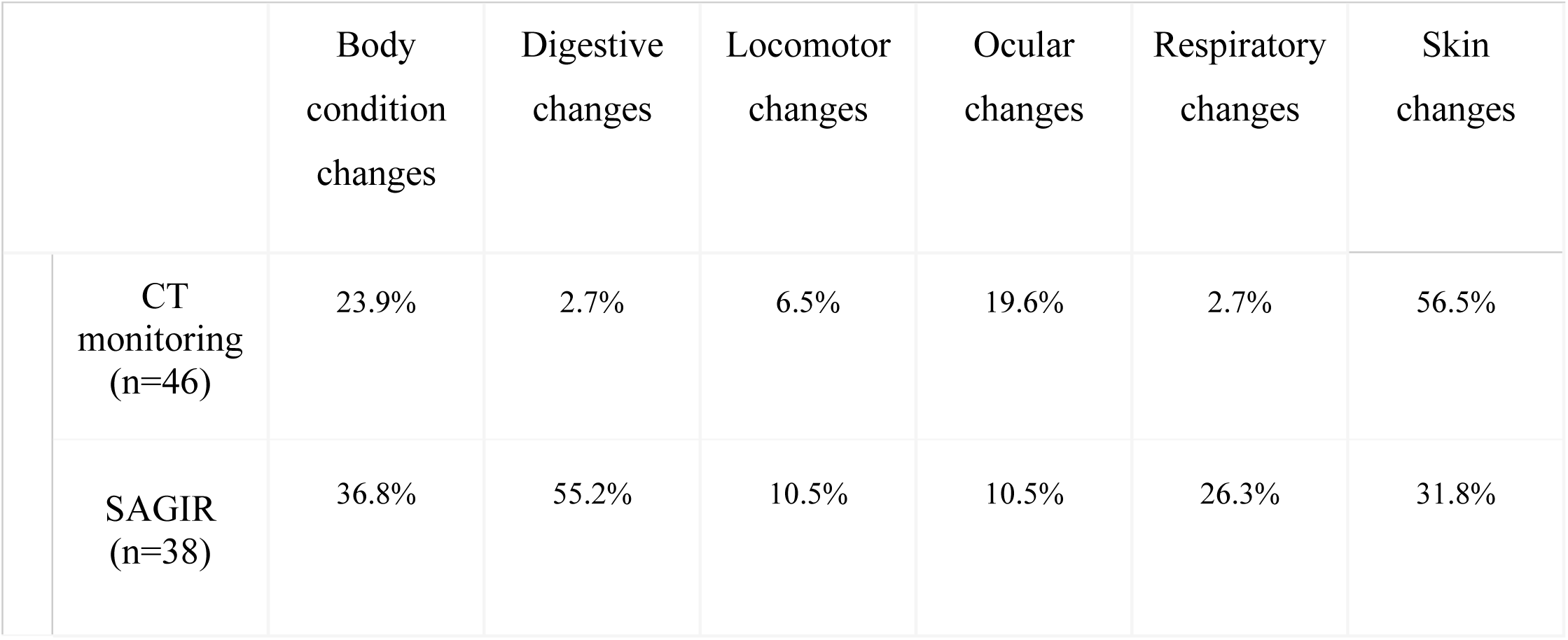
Proportions of lynx with lesions from SAGIR (n=38) and clinical changes from CT monitoring (n=46) in France between 2011 and 2022.

The SAGIR postmortem method detected all lesion categories, in particular digestive, body condition, skin, and respiratory changes and to a lesser extent locomotor and ocular changes (on non-traumatized animals). CT also detects all categories, but preferentially skin, body condition and ocular, and to a lesser extent locomotor, digestive and respiratory changes.

While all disorder categories were detected by both surveillance schemes, in different proportions, some changes were better detected on camera. For example, there was only one case of “rat tail” among the 38 carcasses included here, the disorder accounts for 5% of skin lesions recorded by SAGIR on lynx carcasses, in general. In contrast, over 40% of individuals with skin changes showed a "rat tail" with CT surveillance (n=10). Among the cases of hair loss in necropsied lynx, a severe and fatal case of sarcoptic mange was found in an adult specimen (Figure 7). Extensive hair loss observed in CT surveillance was localized over the abdomen, flanks, medial sides of the legs and tail. It was alternatively visible or absent in recaptured animals suggesting healing or cyclical/seasonal risk. These lesions were not detected through postmortem examination.

**Fig. 7.**
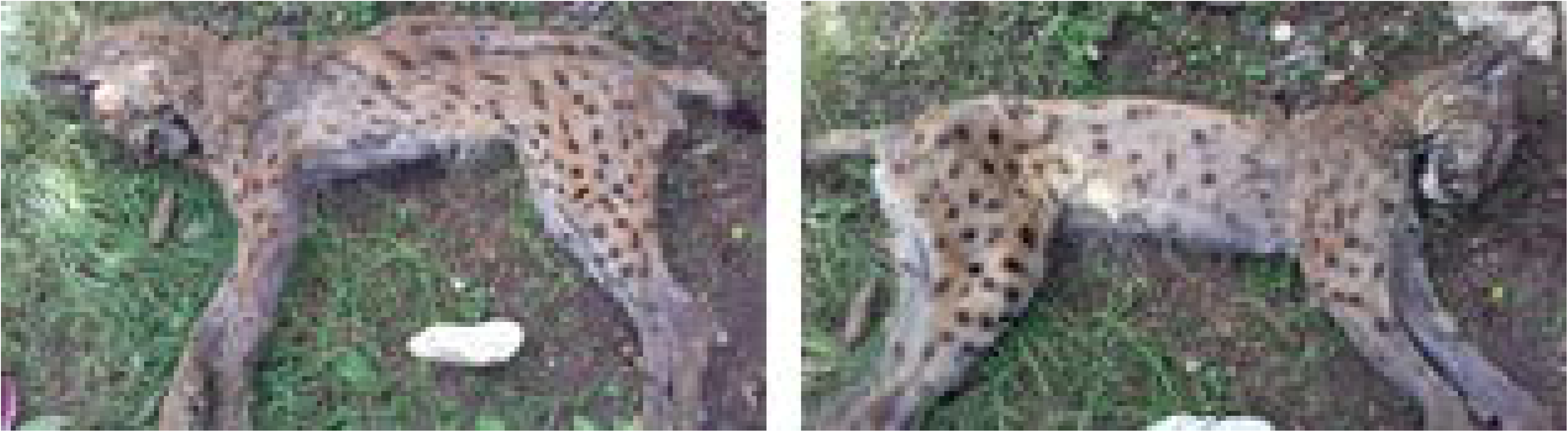
Images of a male lynx carcass at least 9 years old autopsied by the SAGIR network in July 2018, last detected by a camera-trap in February 2017, no changes observed during its lifetime (detected six times between 2011 and 2017) (©OFB25)

Ocular changes observed during necropsies were rare (4/38; 2 individuals with bilateral third eyelid prolapse, one with purulent epiphora and one with a foreign body in the conjunctival sac). CT surveillance detected conjunctivitis and uveitis/keratitis. The congenital absence of the outer ear was exclusively detected by CT, on X individuals, in the year.

## 4. Discussion

### 4.1. Benefits of the method

Opportunistic CT and pictures taken by chance event-based surveillance allows the detection of different health changes in lynx in a far from negligible proportion of individuals (17%) (e.g. skin, body condition, eye, locomotor, behavioral changes, and lack of auriculas). Conversely, the SAGIR postmortem surveillance did not detect changes like uveitis, and severe/cyclic/resolutive hair loss that was not a cause of death. The camera-trap images detected rare cases of absence of auricle, whether this is only a point mutation or linked to inbreeding is currently being clarified. It also offers the possibility of monitoring the evolution of diseases thanks to multiple detections over time as i) lynx can often be identified thanks to their coat pattern, ii) a high proportion of animals were detected multiple times over the monitoring period. CT surveillance is therefore a valuable method for the sanitary surveillance of the Eurasian Lynx in France.

Moreover, the changes detected must be considered as a whole. A variety of disorders in a population can also reflect poor population health linked to causes such as toxic or genetic pressures. There is therefore a different epidemiological significance to the two types of surveillance, and they need to be considered both separately and together.

### 4.2. Differential diagnosis of observed changes

The setup of camera traps, the season (an important parameter for the interpretation of body condition and aspects of the coat), the time of day (day *vs.* night), weather conditions (which modify the perception of colors and contrasts) should be considered, when possible, to interpret the images. To assist biologist’s interpretation, a differential diagnosis is proposed for the main lesions observed (Table 3). To draw up this table, we first looked for each type of change to see if similar cases had been reported in lynx or other wild felids/canids. If no information was available, we then looked for information available for domestic carnivores.

**Table 3:** Differential diagnosis proposed for principal clinical changes observed.

| Type of changes observed | Changes observed | Differential diagnosis | Sources | Comment |
| --- | --- | --- | --- | --- |
| Body condition changes | Distended abdomen | Cardiac congestive failure<br>Hepatopathy<br>Hypoproteinemia<br>Peritonitis<br>Feline infectious peritonitis<br>Neoplasia<br>Hypertension<br>Coagulopathy<br>Gestation<br>Postprandial state | Ettinger et al., (2016) |  |
|  | Thinness | Starvation<br>Lactation period<br>Internal/external parasitosis<br>Chronic viral (ex. FeLV/FIV) or bacterial infection<br>Neoplasia | Ryser-Degiorgis, (2021); Nájera et al., (2019) | Few females were seen very thin in summer |
| Bilateral lack of auriculas |  | Agenesis<br>Traumatic |  | Suspicion of genetic cause (low population heterozygosity), there are others examples of lynx malformation (vascular, skeletal; Ryser-Degiorgis, 2009) |
| Ocular changes | Bilateral prolapse of 3<sup>rd</sup> eyelid | Haw's syndrome<br>Feline dysautonomia | Zwueste & Grahn, (2020); Mendonça et al., (2022)<br>Hernandez et al., (2010); Ketrings et al., (2012) | Described mainly in young cats with diarrhea, no definite known physiopathology (hypothesis of viral or parasitic causes)<br>Central neurological manifestation of intestinal damage, often associated with diarrhea |
|  | Conjunctivitis | Infection à *Chlamydia felis*<br>Others viral or bacterial infections | Marti et al., (2019) | Common in domestic cats, Detected in a lynx in Switzerland |
|  | Uveitis/keratitis | Traumatic<br>Infectious (viral, bacterial, fungus)<br>Tumoral | Ketring et al., (2012) | Neovascularisation during cicatrisation visible on few images (supplementary materials) |
| Skin changes | Wounds/scars | Traumatic |  |  |
|  | Symmetrical hair loss | High probability of non-inflammatory cause:<br>- Endocrine (pathology sexual dysendocrinia, hypothyroidism, Cushing syndrome or an equivalent of “Canine Flank Depilation”<br>- Genetic (Follicular and seasonal dysplasia)<br>- Toxic<br>- Traumatic<br>- Autoimmune<br>- Behavioral<br>- “Rat tail” syndrome<br>- Moults (physiological) | Coyner, (2019)<br>Welle et al., (2023);<br>Verschuuren, (2022)<br>Gordon et al., (2023) | Possibly associated with others metabolic changes<br>Known phenomenon in squirrels<br>Observed in captive wild felids (Thorel et al., 2020; Napier et al., 2018)<br>Unknown aetiology but already observed in Switzerland (Ryser-Degiorgis, 2020) |
|  | Asymmetrical hair loss | High probability of inflammatory cause:<br>- Sarcoptique or notoédrique mange<br>- Extensive depilation cause by pulicose<br>- Fungal<br>- Bacterial<br>- Neoplasia<br>- *Otodectes Cynotis* infection | Moraillon & Legeay, (2004);<br>Paterson, (2016);<br>Coyner, (2019)<br>Ryser-Degiorgis, (2002); Schmidt-Posthaus et al., (2002)<br>Degiorgis et al., (2001) | Found on a very emaciated and depilated dead lynx at histology. Very common in Europe mammal population, threat for ungulates and wild carnivores especially foxes (lynx prey) (Escobar, 2021; Pisano et al., 2019). Suspected on 1 dead individual at histology |

However, we cannot make consistent parallels, as we have only 5 identified lynx that have been observed by CT then registered and necropsied by SAGIR. Moreover, we have few histological samples analyzed in the population. To carry out a more precise study, a few individuals should be equipped with GPS to be necropsied by the SAGIR network if they were to die and to have a histological diagnosis of unknown lesions and a better representation of cause of death. Lynx admitted to wildlife rescue centers could provide an opportunity as well to reach a more precise diagnosis (cyclic hair loss for example).

### 4.3. Biases of this method

The main, obvious limit of camera trap surveillance is that it only detects diseases that lead to external signs. Furthermore, as it is opportunistic, observation pressure was not homogeneous over time and space and cannot be precisely known over the course of the study, as observers and protocols were diverse. Thus, the frequency of observation of a clinical change is neither an accurate nor a precise estimator of prevalence. Moreover, numerous factors influenced the quality of data. Blin et al. (unpublished results) showed on this dataset that the detection of skin changes is easier during the night, whereas body condition changes are better detected during the day. Blurs in the picture, the presence of a flash, the shooting angle, and the percentage of the individual visible in the picture were also critical to detect or not a clinical change with a camera trap image.

The high heterogeneity of the data is both a strength and a weakness for earlier detection and characterization of clinical changes in lynx in comparison with SAGIR surveillance. It is a strength because of the quantity of data obtained and good detection of some changes that better visible in a specific kind of setup, such as eye damage with a flash at night. But it is also a weakness because the collection of data is not standardized. Trends can be detected, however, the photographic equipment used before 2010 was not the same quality as that available today, and the number of CTs used for ecological monitoring was much lower. It is therefore difficult to state whether certain changes were really emerging, or whether they were not detected/detectable with the technical means available before 2010. When the quality of images was high enough, we could even detect small scars or ocular damage such as conjunctivitis or uveitis. As previously described for sarcoptic mange by Olega et al. (2011), severe changes (e.g. here severe hair loss, cachectic body condition or auricular lack) were detected more easily and with a higher level of confidence than mild to moderate lesions. After the first screening step, 17% of individuals had at least one image with suspected clinical changes, but 37% initially flagged by the student were downgraded to “No visible changes” by the expert veterinarian. Moreover, some types of change are more difficult to confirm: minimal hair loss, “rat tail.,” when a reflection of the flash on the coat can cast doubt on depilation; a lack of sharpness on a moving lynx making it difficult to assess its body condition; too large a distance between the animal and the camera limiting the characterization of skin or eye lesions. For this purpose, a second, complementary study has been carried out in parallel. It has characterized the factors that interfere with the detectability of changes, by establishing quality criteria for the observation of photographs taken by CT as part of the health surveillance of the Eurasian lynx in France (Blin et al., unpublished results). This shows the importance of multiplying the number of qualified observers and the number of observations for each data. Indeed, we note that during our study the observer’s eye is gradually trained as the observations are carried out. Thus we publish an atlas of normal (physiological) images and another with images of pathological changes, alongside this paper, to accelerate and standardize the learning process to screen data for each observer.

### 4.4. The relevance of multimodal monitoring

As SAGIR and CT syndromic surveillance are event-based surveillance, the patterns identified in the data are driven both by the observation process and the underlying biological process of interest (‘state process’) (Chadwick et al. 2023) and it is therefore difficult to differentiate between the contribution of each process. Observation process is for example affected by the detectability of carcasses or live lynx (ex. carcasses resulting from collisions have an almost 100% chance of being spotted, unlike most of the carcasses of individuals that have died in remote areas). The rate of reporting to RLL or SAGIR, the state of preservation of the carcass or the quality of images all result in heterogeneous observation pressure in time and space. Multiplying surveillance modalities could improve the understanding of the state process (i.e. the factors influencing the animal’s true health status or cause of death). Our results suggest that carcass-based and CT surveillance are complementary, the latter detecting some changes earlier than SAGIR, which primary focus is cause of death. Some skin and ocular changes are difficult to detect by SAGIR because they are mostly sublethal and the probability to find them in animals dead from other causes is low. Furthermore, eyes decay rapidly after death so pathological signs are difficult to detect. Conversely, CT syndromic surveillance allows only a rough description and making some diagnostic hypotheses. Postmortem or clinical examination on similar cases admitted to rehabilitation centers must complement the diagnosis to count actual cases of disease. A rise in abnormal signs on camera images could prompt further examination in necropsied animals to detect infectious diseases or toxic compounds for example. This study has shown that different types of changes can be detected by CT and that the more surveillance modalities available, the better the detection. However, even if multiplying the number of CT performed seems an interesting way of increasing the amount of data obtained, it is not possible indefinitely, and in particular if the setup requires two cameras per site. Health monitoring with camera traps should be seen as a by-product of lynx monitoring for demographic aims. This, associated with a more systematic comparison of lesions observed on carcasses previously seen alive by CT, would give us a better chance of making a diagnosis of certainty in case of a sharp increase in pathological visible changes, with the support of additional examinations on carcasses (histology, bacteriology, PCR, mycology, etc.).

### 4.5. Ethical context of CT wildlife surveillance

To put this work into perspective, our attention was drawn to the ethical aspect of CT installations in the wild. In fact, the greater the number of cameras installed, the greater the disturbance can be, caused by the presence of humans during placement and retrieval, particularly in remote areas. However, camera traps are set along trails and forest roads and are mainly checked during the day. Hence this process has a low probability to be a significative disturbance in a multi-use landscape such as the Jura and Alps.

Moreover, during CT monitoring, images or videos of humans are regularly taken unintentionally. These data can be potentially harmful to people who are filmed without their knowledge (Franchini et al, 2022). Some institutions already use artificial intelligence to remove empty pictures and human activities to save time and increase the level of privacy guarantees. The OFB always uses posters associated with CT to explain the role of these cameras. However, research teams generally limit large-scale communication about the presence of cameras, for fear of vandalism. Also, recommendations are being issued on the processing of unintentionally collected human data, mainly to deal with cases where illegal acts that may be filmed by these cameras (Sharma et al., 2020).

## 5. Global perspectives and Conclusion

Epidemiological surveillance of wildlife diseases is a challenge, especially when it comes to elusive or rare species. Improvement is needed: for example, data are scarce, with many biases, biological samples are of varying quality, diagnosis is adapted from domestic species, stakeholders working on the species are in little or no contact with health-related studies, etc. The CT opportunistic syndromic surveillance proposed in this study, driven by a network of field observers, is a way to reinforce lynx disease surveillance, as it allows to detect changes of interest for the population and monitor the clinical evolution of a change (multiple detections). This non-invasive method can be adapted to other species, after establishing suitable frames of reference. It is of particular advantage on individually identifiable species, such as most felids but also painted dogs, hyenas, giraffes or zebras.

Finally, this method complements other surveillance systems and calls for transdisciplinary collaboration. However, the setup and workflow of this method of collecting data requires interoperability between surveillance systems, to ensure traceability between individuals (living and dead), training sessions of observers to better detect changes with help of training images and additional human resources. This method represents a low material cost as it makes use of images collected for biological monitoring, but it is time-intensive, because of the initial screening, annotation by field observers, followed by expert validation of the images. Artificial intelligence tools could help to improve all three of these steps and eventually reduce the manpower requirements.

## Data availability statement

The raw data supporting the conclusions of this article will be made available by the authors, under specific conditions (because it includes confidential data for the conservation of the species). Please contact the corresponding author.

## Supplementary material

1. Atlas for the observation of the Boreal lynx by camera trapping - Apparent health status
2. Atlas for the observation of the Boreal lynx by camera trapping - Observable changes
3. Complete protocol for the clinical observation of boreal lynx by camera trap

## Author contributions

LL: Writing – original draft, conceptualization, methodology, project administration, formal analysis, visualization. DC: Writing – Review & editing, conceptualization, methodology, project administration, supervision, data curation, LB: Writing – review & editing, conceptualization, methodology, data curation SBa: Writing – Review & editing, conceptualization, methodology, project administration, supervision, formal analysis. SBo, FZ, NT, KL: Writing – Review & editing, M-PR-D: Conceptualization, methodology. PG: Writing – Review & editing, conceptualization, methodology, project administration, supervision, formal analysis, AD: Writing – Review & editing, conceptualization, methodology, project administration, supervision, formal analysis.

## Acknowledgements

We would like to thank all contributors to this work: the Loup-Lynx network and SAGIR teams, the veterinary departmental laboratories.

## Funding resources

This research did not receive any specific grant from funding agencies in the public, commercial, or not-for-profit sectors.

