## Supplemental_materials for "What if wildlife health surveillance was not just for veterinarians? - Opportunistic use of population monitoring by camera traps for syndromic surveillance of the Eurasian lynx (*Lynx lynx*)"

### Supplemental materials 1 and 2: Atlases

The Atlas of apparent health status SM1 and the Atlas of observable changes SM2 that are referenced in the manuscript will not be published as a preprint, pending publication in a peer-reviewed journal, however, the authors may provide them upon reasonable request.

### Supplemental material 3: Complete protocol for the clinical observation of boreal lynx by camera

The aim of this document is to provide a precise description of the analysis method to be used for health monitoring by CT of Boreal Lynx in France.

1.1. Initial training

Watch the two photographic reference tools for observing the Boreal Lynx by CT surveillance named "State of apparent good health", to train your eye on lynx in good health and to learn about the difficulties encountered when viewing images and videos for health surveillance. Then look at the Boreal Lynx photographic reference tool named "Observable changes" to get an idea of the different types of changes encountered and the key points to check when analysing each image or video.

1.2. Clinical observation protocol for lynx health surveillance:

This is the list of elements to look for in each image or video. Of course, not all points can be observed on every file, but this will ensure that you do not overlook an element when viewing. First, it is important to take into account the context of the capture. The season is an important parameter to know to observe the animal's body condition and coat. This information is directly accessible on the image or video, as the date is visible. In addition, you need to consider the time of shooting, day or night, as well as weather conditions, which modify the perception of colors and contrasts.

1.2.1. Clinical observation applied to the photographic medium:
- Body condition (protruding ribs, protruding transverse apophyses (= bony tips of vertebrae), protruding hip and buttock tips)
- General condition
- Posture (head carriage, back carriage, etc.)
- Behavior/Neurological changes (abnormally protruding tongue, exaggerated locomotor movements, prostration, absence of escape, etc.)
- Head:
  - Jaw (fracture, prognathism (= lower jaw advanced in relation to upper jaw), appearance of teeth)
  - Eyes (procidence 3rd eyelid, ulcer, exophthalmos/enophthalmos (= eyeball protruding/inserted), discharge, closed eye, white eye, red eye (inflammation) etc.). A specific method for optimizing the characterization of ocular lesions was set up with the help of an ophthalmology specialist: use of Photoshop® software to contrast and zoom in on the eyes and visualize the eye fundus.
  - Nose (hyperkeratosis (= increased skin thickness), nasal discharge, etc.)
  - Ears (scabs, lesions, ear shape, etc.)
- Digestive system: Anal staining or lack of hair on hocks (may indicate chronic diarrhea), prolapse (= externalization of rectum), hypersalivation, bloating.
- Cutaneous system (head to tail): alopecia, mass, trauma, crusting, subcutaneous edema, etc.
- Respiratory system: panting.
- Musculoskeletal system: fracture, joints, plumbness, amyotrophy, loss of limb, etc.
  1.2.2. Clinical observation applied to video support:
- Body condition (protruding ribs, protruding transverse processes, protruding hip and buttock tips)
- General condition (good or suspect (e.g. shaggy hair))
- Posture (head carriage, back carriage, etc.)
- Behavior/Neurological changes (abnormally protruding tongue, exaggerated locomotor gestures, groping, ataxia (= coordination change), prostration, failure to flee from man, tremors, pruritus, circling, vocalization, etc.).
- Head:
  - Jaw (fracture, prognathism, appearance of teeth)
  - Eyes (procidence 3rd eyelid, ulcer, exophthalmos/enophthalmos, discharge, closed eye, white eye, red eye, etc.) A specific method for optimizing the characterization of ocular lesions was set up with the help of an ophthalmology specialist: use of Photoshop® software to contrast and zoom in on the eyes and visualize the eye fundus.
  - Nose (hyperkeratosis, nasal discharge, etc.)
  - Ears (scabs, lesions, ear shape, etc.)
- Digestive system: Anal staining or lack of hair on hock (may indicate chronic diarrhea), prolapse, hypersalivation, bloating, vomiting.
- Cutaneous system (head to tail): alopecia, mass, trauma, scabs, edema, etc.
- Respiratory system: panting, respiratory frequency, sneezing, etc.
- Musculoskeletal system: fracture, joints, plumbness, amyotrophy, loss of limb, lameness, locomotor changes, etc.
  1.3. Recording observed changes and updating the database

In the file listing all the events recorded in the database, three columns are added: the first to indicate the type of change (e.g. "Tskin"), the second to specify the nature of the change (e.g. " Tskin_Severe_Alopecia ") and the last to specify the name(s) of the file(s) where the change is visible to the observer in the series of photos/videos of the event concerned. Finally, the comment column could be used for any comments that seem necessary to describe the change by the biologist.
